# A pathogenicity locus of *Streptococcus gallolyticus* subspecies *gallolyticus*

**DOI:** 10.1101/2022.03.15.484266

**Authors:** John Culver Taylor, Ritesh Kumar, Juan Xu, Yi Xu

## Abstract

*Streptococcus gallolyticus* subspecies *gallolyticus* (*Sgg*) is known to be strongly associated with colorectal cancer (CRC). Recent functional studies further demonstrated that *Sgg* actively stimulates CRC cell proliferation and promotes the development of colon tumors. However, the *Sgg* factors important for the pro-proliferative and pro-tumor activities of *Sgg* remain unclear. Here, we identified a chromosomal locus in *Sgg* strain TX20005. Deletion of this locus significantly reduced *Sgg* adherence to CRC cells and abrogated the ability of *Sgg* to stimulate CRC cell proliferation. Thus, we designate this locus as the *<u>S</u>gg* <u>p</u>athogenicity-<u>a</u>ssociated <u>r</u>egion (*SPAR*). More importantly, we found that *SPAR* is important for *Sgg* pathogenicity *in vivo*. In a gut colonization model, mice exposed to the *SPAR* deletion mutant showed significantly reduced *Sgg* load in the colonic tissues and fecal materials, suggesting that *SPAR* contributes to the colonization capacity of *Sgg*. In a mouse model of CRC, deletion of *SPAR* abolished the ability of *Sgg* to promote the development of colon tumors growth. Taken together, these results highlight *SPAR* as a critical pathogenicity determinant of *Sgg*.

## Introduction

*Streptococcus gallolyticus* subspecies *gallolyticus* (*Sgg*) is a group D streptococcal bacterium, and an opportunistic pathogen belonging to the *Streptococcus bovis/Streptococcus equinis* complex (SBSEC) (1). *Sgg* causes bacteremia and infective endocarditis (IE), and has a long-standing clinical association with CRC, supported by extensive studies and case reports dating back to the 1950s (2–15). Patients with *Sgg* IE and/or bacteremia have concomitant colon adenomas or adenocarcinomas at a much higher rate (~60%) than that of the general population (8). Moreover, a prospective study found that 45% of patients with *Sgg* IE developed colonic neoplastic lesions within 5 years of IE diagnosis compared to 21% of patients with IE due to closely-related enterococci, further supporting the association of *Sgg* with CRC (6). In CRC patients with no symptoms of IE or bacteremia, *Sgg* was found to preferentially associate with tumor tissues compared to normal adjacent tissues or normal tissues in multiple studies around the world (4, 16–19).

Functionally, *Sgg* stimulates the proliferation of human CRC cells in a β-catenin dependent manner *in vitro* and in a xenograft model (4). Exposure to *Sgg* also led to larger and more advanced tumors in an azoxymethane (AOM)-induced CRC model and in a colitis-associated CRC model (4, 15). These results indicate that *Sgg* possesses pro-proliferation and pro-tumor activities. Previous work also indicated that there are variations among *Sgg* strains in their ability to stimulate human CRC cell proliferation. Some *Sgg* strains such as ATCC43143 showed significantly reduced capacity to adhere to HT29 cells, to stimulate the proliferation of these cells and to promote tumor development *in vivo* (2), suggesting that there are specific *Sgg* factors contributing to the pro-proliferative and pro-tumor activities of *Sgg*. The identity of these specific *Sgg* factors was unknown.

Here, we describe work characterizing a chromosomal locus of *Sgg* strain TX20005. Deletion of this locus significantly impaired the ability of *Sgg* to adhere to cultured CRC cells and abrogated the ability of *Sgg* to stimulate CRC cell proliferation. *In vivo*, the deletion mutant displayed significantly reduced capacity to colonize normal mouse colons. More importantly, the mutant lost the ability to promote the development of colon tumors. Given the role of this locus in *Sgg* pathogenesis, we have coined this locus as the *Sgg* pathogenicity-associated region (*SPAR*).

## Materials and Methods

### Bacterial strains, cell lines and growth conditions

*Sgg* strains were routinely grown in brain heart infusion (BHI) broth or tryptic soy broth at 37°C with shaking, or on BHI or TSB agar plates at 37°C overnight (Teknova). For co-culture experiments and animal studies, stationary phase bacterial cultures were pelleted, washed with phosphate-buffered saline (PBS), pH 7.4, resuspended in PBS containing 15% glycerol, aliquoted and stored at −80°C. Aliquoted stocks were thawed, washed with PBS, diluted in appropriate media to obtain the desired concentration and directly added to cells or administered to mice. Human colon cancer cell lines HT29, HCT116, and SW480, as well as the HEK293 cell line were grown in Dulbecco’s Modified Eagle’s Medium F-12 50/50 (DMEM/F-12, GIBCO) supplemented with 10% FBS at 37°C with 5% CO_2_ in a humidified chamber. Cells from less than 30 passages were used in the experiments.

### Preparation of culture supernatants (CSs)

Supernatants from stationary phase *Sgg* cultured in BHI were filtered-sterilized through a 0.2μm filter and concentrated approximately 10 – 20-fold using centrifugal concentrators (3kD molecular weight cut off). The concentrated CSs were aliquoted and stored at −80°C. A vial is thawed and diluted in the appropriate media for adherence or cell proliferation assays and used immediately.

### Adherence assay

This was performed as described previously (3). Briefly, HT29 cells were seeded at a density of 1×10^6^ cells/well in a 24 well plate and allowed to attach overnight. Cells were incubated with bacteria at a multiplicity of infection (MOI) of 10 in the absence or presence of CSs (diluted to 1x) for 1 hour at 37°C with 5% CO_2_. Cells were washed thrice in PBS to remove unattached bacteria, lysed with 1mL of 0.01% Triton X-100 (Sigma) and dilution plated onto BHI or TSB agar plates. Adherence was calculated as the percentage of adhered bacteria vs. total bacteria added.

### Cell proliferation assay

This was performed as described previously (3) with slight modifications. Briefly, cells were seeded in 96 well plates at a concentration of ~1×10^4^ cells/well and incubated overnight. Cells were then incubated in fresh DMEM/F-12 containing 10% FBS in the absence or presence of *Sgg* bacteria (MOI = 1) or *Sgg* CSs (1x) for a total of 24 hours. Trimethoprim was added after 6 hours of incubation (1 μg/mL final concentration) to inhibit bacterial growth, as previously described. The number of viable cells was determined using the cell counting kit (CCK)-8 kit following the instructions of the supplier (Apex Bio).

### Western Blot

Detection of β-catenin and PCNA was carried out as described previously (3). Briefly, HT29 cells were seeded at a density of 1×10^6^ cells per well in 6-well plates and incubated overnight. The cells were then incubated with media only or media containing bacteria (MOI=1) for 9 hours. Total cell lysates were subjected to SDS-PAGE, transferred, and probed with antibodies against β-catenin (1:1000), PCNA (1:1000), and β-actin (1:1000) (Cell Signaling Technology), followed by incubation with HRP-conjugated secondary antibodies (1:3000). Band intensities were measured using FIJI ImageJ and normalized to those of β-actin.

### Deletion of *SPAR*

This was performed following a procedure described previously (3). Briefly, the ~ 1kb region upstream of *sparA* and the ~1kb region downstream of *sparL* were synthesized by Genscript and cloned into pUC57 (Genscript). The insert was then subcloned into a temperature sensitive conjugative plasmid pG1-*oriT*_TnGBS1_. The insert sequence was verified by Sanger sequencing. The construct was introduced into *S*. *agalactiae* NEM316 by electroporation and then into *Sgg* strain TX20005 by conjugation under the permissive temperature. PCR was used to screen for double cross-over deletion mutants. Deletion was further confirmed by PCR amplification of the regions spanning the deleted fragment and DNA sequencing of the PCR product.

### RNA extraction, cDNA synthesis and PCR

This was performed following the method described previously (3). Briefly, RNA was extracted from TX20005 cultured in the presence of HT29, as well as from the colonic tissues of mice that had been orally gavaged with TX20005. RNA was treated with DNase and synthesized using a ProtoScript II First Strand cDNA Synthesis Kit (NEB). Primers used in PCR amplification are: *sparA*, forward 5’GCAAGCTGGTCGAACAGAAC and reverse 5’GCTTCTATGGTTGGGGCTAGA; *sparD*, forward 5’ GGAGGTGGATCCAACAAGGG and reverse 5’CAGGTTCCTCGATAGCCAGC; and *sparG*, forward 5’TCAGTTGTTAGCGGATGCGT and reverse 5’ CCCTTTATTGCTTGTGCTCCC.

### Growth curves

Overnight cultures of *Sgg* were inoculated into fresh BHI broth at 1:100 dilution and grown at 37°C with shaking. Samples were taken at 0, 3, 6, 9, 12, and 24 hours, dilution plated onto agar plates, incubated for 24 hours and colonies enumerated.

### Animal experiments

Animal studies were performed in accordance with protocols approved by the Institutional Animal Care and Use Committee at the Texas A&M Health Science Center, Institute of Biosciences and Technology. Mice were fed with standard ProLab IsoPro RMH3000 (LabDiet). **1) Colonization.** This was performed as previously described (3) with slight modifications. Briefly, 6-week-old A/J mice, sex matched (Jackson Laboratory), were treated with ampicillin at a concentration of 1 g/L in drinking water for 3 days and switched to antibiotic-free water 24 hours prior to the administration of bacteria. *Sgg* was orally gavaged at a dose of ~1 x 10^9^ CFU/mouse. Colons and fecal materials were collected at day 1, 3, and 7 post-gavage. Samples were weighed, homogenized in sterile PBS in a TissueLyser (Qiagen), dilution plated onto Enterococcus Selective Agar (ESA) plates and incubated at 37°C for 24–48 hours to enumerate *Sgg* colonies. **2) AOM-induced model of CRC.** This was performed as previously described (3) with slight modifications. Briefly, 6-week-old A/J mice, sex matched were treated with 4 weekly i.p. injections of AOM (10 mg/kg body weight), followed by ampicillin in drinking water (1 g/L) for 7 days and switched to antibiotic-free water 24 hours prior to the first oral gavage with *Sgg*. Mice were orally gavaged with bacteria at ~ 1×10^9^ CFU/mouse or saline (n = 11-12 per group) once per week for 12 weeks. Mice were euthanized one week after the last oral gavage by CO_2_ inhalation followed by cervical dislocation. Colon and fecal pellets were collected. The number and size of macroscopic were recorded. Visual evaluation of colons was carried out by a blinded observer. A random subset of fecal pellets were weighed, homogenized in sterile PBS and dilution plated onto ESA plates.

### Statistical analysis

GraphPad Prism 9 was used for statistical analyses. Two-tailed unpaired *t*-test was used for pairwise comparisons to assess the significance of differences between two groups in cell proliferation assays, western blot analysis, and adherence assays. The non-parametric Mann-Whitney test was used to assess the significance of differences of results between groups in animal studies. Two-way ANOVA was used to analyze the bacterial growth curves. Ns, not significant; *, *p* < 0.05; **, *p* < 0.01; ***, *p* < 0.001; ****, *p* < 0.0001.

### Ethics statement

Animal studies were performed in accordance with protocols (IACUC#2017-0420-IBT) approved by the Institutional Animal Care and Use Committee at the Texas A&M Health Science Center, Institute of Biosciences and Technology. The Texas A&M University Health Science Center—Institute of Biosciences and Technology is registered with the Office of Laboratory Animal Welfare per Assurance A4012-01. It is guided by the PHS Policy on Human Care and Use of Laboratory Animals (Policy), as well as all applicable provisions of the Animal Welfare Act.

## Results

### The *Sgg SPAR* locus

We compared the genome sequence of TX20005 (NZ_CP077423.1) with that of ATCC 43143 (NC_017576.1), an *Sgg* strain that is defective in stimulating CRC cell proliferation or promoting the development of colon tumors, using the multiple genome alignment tool MAUVE (20). The SPAR locus is one of the regions in the chromosome of TX20005 that display differences from ATCC 43143. The locus is comprised of 12 genes (*sparA* to *sparL*) (Fig. 1A). The protein sequences encoded by these genes were analyzed for secretion signals using SignalP - 6.0 (21). None of them contains typical secretion signals recognized by the Sec or Tat translocon. SparA, SparK and SparL are predicted to contain transmembrane helices (TMHMM2.0) (22) and are thus putative transmembrane proteins. Homology search using Protein BLAST showed that SparA to C are hypothetical proteins of unknown function (Table 1). SparD to SparF exhibit homology to predicted ATP-dependent endonuclease of the OLD family, GIY-YIG endonuclease/PcrA/UvrD helicase, and GntR family transcriptional regulator, respectively. SparG to SparJ display features of effectors secreted by the type VII secretion system (T7SS). SparG, H and J are homologous to T7SS effectors, LXG domain-containing protein and T7SS effector LXG polymorphic toxin. SparG, H and I are of 91, 102 and 137 amino acids in length, in keeping with T7SS effectors belonging to the WXG100 family. Analysis using HHpred (23) showed that SparG and SparI fold into a four-helical bundle structure typical of ESAT-6-like WXG100 proteins, subfamily of T7SS effectors, with a probability of 96.48% and 97.97%, respectively. Spar H, I and J also contain a conserved sequence motif HxxxD/ExxhxxxH (H denotes highly conserved and h less conserved hydrophobic residues) that is considered to be important for the secretion of WXG100 proteins by T7SS (24) (Supplemental Fig. S1). Immediately downstream of the *sparJ* gene are two genes that encode putative TipC family immunity proteins. Genes encoding T7SS effector LXG polymorphic toxins are typically clustered together with genes encoding their cognate immunity proteins (25–27). The arrangement of *sparJ*, *K* and *L* further suggests that SparJ is a putative T7SS effector LXG polymorphic toxin and SparK and L are likely the cognate immunity proteins.

**Fig 1.**
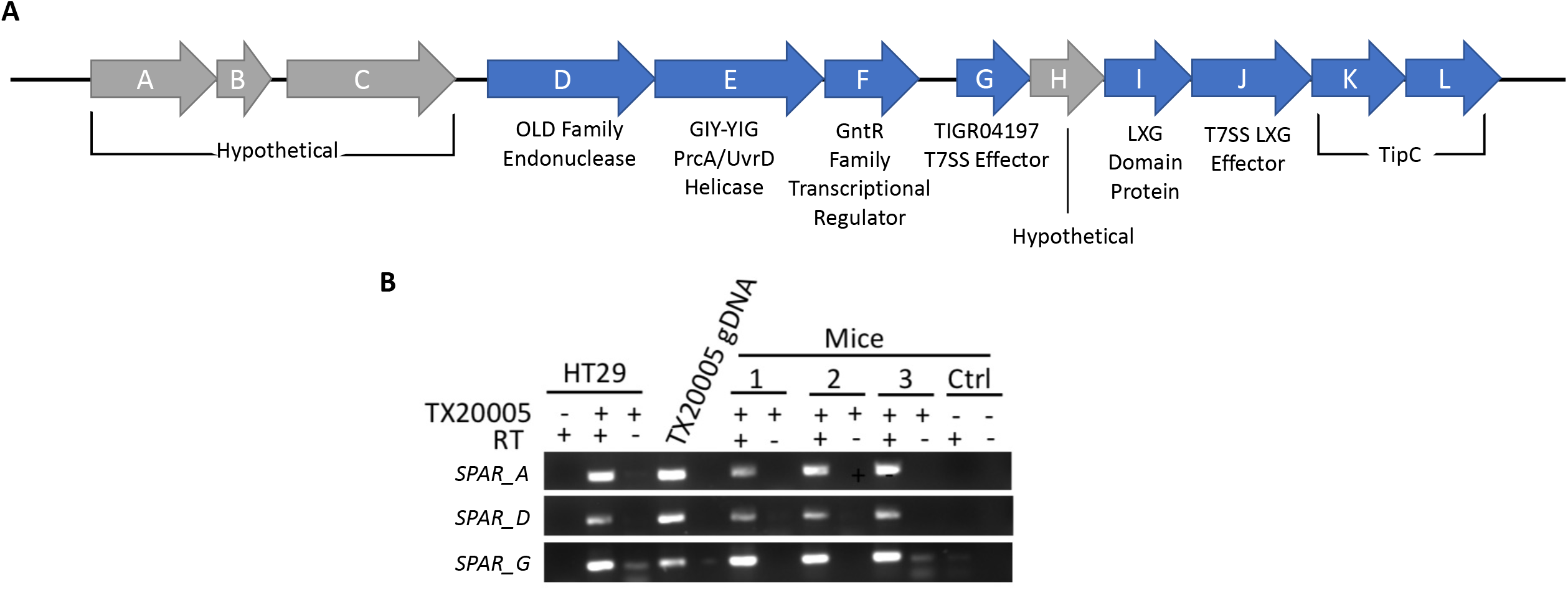
*SPAR* genetic organization and expression. **A.** The *SPAR* locus. Genes encoding proteins with homology to proteins of known functions or containing characteristics of certain protein families are colored in blue, while genes encoding hypothetical proteins of unknown function are colored in gray. **B. RT-PCR.** cDNA synthesized from RNA extracted from HT29 cells co-cultured with TX20005 and the colonic tissues of three mice that had been orally gavaged with TX20005 were used as a template for the RT-PCR. cDNA from HT29 cells only and from mice orally gavaged with saline were used as controls to show the specificity of the PCR primers. RNA samples that had not been treated with reverse transcriptase (RT) were used as controls to show the lack of DNA contamination.

**Table 1.** Proteins encoded by the *SPAR* genes.

| Gene | Length (aa)<sup>a</sup> | Predicted Gene Function<sup>b</sup> |
| --- | --- | --- |
| *sparA* | 424 | Hypothetical Protein |
| *sparB* | 53 | Hypothetical Protein |
| *sparC* | 582 | Hypothetical Protein |
| *sparD* | 656 | OLD-Family Endonuclease |
| *sparE* | 609 | GIY-YIG Endonuclease/PcrA/UvrD Helicase |
| *sparF* | 232 | GntR-Family Transcriptional Regulator |
| *sparG* | 91 | TIGR04197-Family T7SS Effector |
| *sparH* | 102 | Hypothetical Protein |
| *sparI* | 137 | LXG domain-containing protein |
| *sparJ* | 417 | T7SS effector LXG polymorphic toxin |
| *sparK* | 199 | TipC Family Immunity Protein |
| *sparL* | 197 | TipC Family Immunity Protein |

<sup>a</sup>Based on the annotated genome of TX20005 (NZ\_CP077423.1)
<sup>b</sup>The amino acid sequence of each protein was used to search the non-redundant protein
sequences database (updated on 04/05/2022) at NCBI using Protein BLAST.

We next tested if the *SPAR* genes were expressed *in vitro* and *in vivo* by RT-PCR (Fig. 1B). We focused on the expression of *sparA*, *D* and *G*, under conditions that are relevant to the pro-proliferation and pro-tumor activity of *Sgg*. All three genes were expressed when the bacteria were co-cultured with HT29 cells and in the colonic tissues collected from mice orally gavaged with TX20005 (Fig. 1B).

### *SPAR* is important for *Sgg* adherence

To investigate the role of the *SPAR* locus in the pathogenic potential of *Sgg*, we generated a mutant in the TX20005 background, in which all 12 genes were deleted (TX20005Δ*SPAR*). We verified that the deletion does not affect bacterial growth (Fig. 2A). We sought to determine if deletion of the *SPAR* locus had any effect on the adherence capacity of *Sgg*. The results showed that the mutant strain exhibited a significantly reduced adherence to HT29 cells compared to the wild type (WT) parent strain TX20005 (~2% vs ~9%) (Fig. 2B), suggesting that the *SPAR* locus is important for *Sgg* adherence to host cells. Previous work showed that T7SS-secreted factors of *Sgg* significantly enhanced the adherence capacity of the WT bacteria (3). We tested whether filter sterilized culture supernatants (CS) from TX20005Δ*SPAR* can enhance the adherence capacity of TX20005. Addition of CS from the WT TX20005 significantly enhanced the adherence of WT bacteria, whereas addition of the mutant CS did not cause any increase in the adherence of WT bacteria (Fig. 2B, indicated by the arrow), suggesting that *SPAR*-dependent secreted factors contribute to *Sgg* adherence.

**Fig 2.**
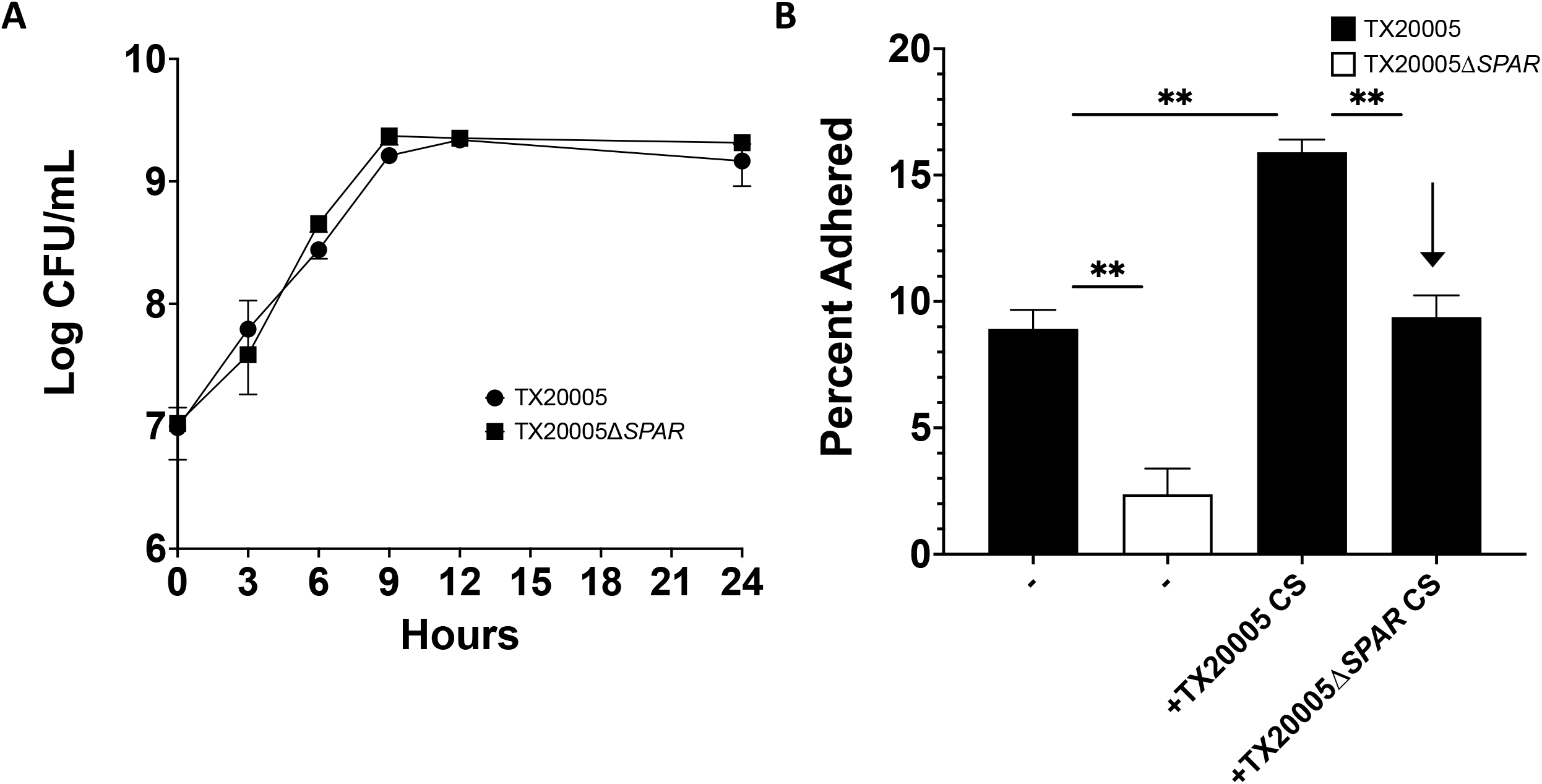
SPAR is important for *Sgg* adherence but does not impact growth. **A. Growth Curve.** A single colony of WT or mutant bacteria was inoculated into 1mL of fresh BHI and incubated at 37°C overnight with shaking. The overnight culture was then diluted 1:100 with fresh BHI and incubated at 37°C with shaking. 100μL of the culture were taken at the 0, 3, 6, 9, 12, and 24 hours for serial dilution and plating. Results were combined from three biological replicates. **B. Adherence**. HT29 cells were seeded at a density of 1×10^6^ cells per well and incubated with TX20005 or TX20005*ΔSPAR* (MOI=10) in media only or in the presence of CS from TX20005 or TX20005*ΔSPAR* for 1 hour. Percentage adherence was calculated as the percentage of adhered bacteria vs total bacteria added. Results were combined from three biological replicates. Unpaired two-tailed *t* tests were used. **, *p*<0.01; NS, not significant.

### *SPAR* is required for *Sgg* to stimulate CRC cell proliferation

*Sgg* has been previously shown to stimulate cell proliferation in human CRC cell lines HT29 and HCT116, but not in SW480 or HEK293 cell lines (2–4). We examined the ability of TX20005Δ*SPAR* to stimulate cell proliferation in these cell lines. HT29, HCT116, SW480, and HEK293 cells were co-cultured with TX20005 or TX20005Δ*SPAR*. As anticipated, the WT strain significantly stimulated HT29 and HCT116 cell proliferation compared to untreated cells (Fig. 3A and 3B). In contrast, HT29 and HCT116 cells co-cultured with the mutant did not exhibit any significant increase in cell proliferation, indicating that *SPAR* is required for *Sgg*-stimulated CRC cell proliferation. Neither the WT nor the mutant had any effect on the proliferation of SW480 or HEK293 cells (Fig. 3C and 3D), as expected.

**Fig 3.**
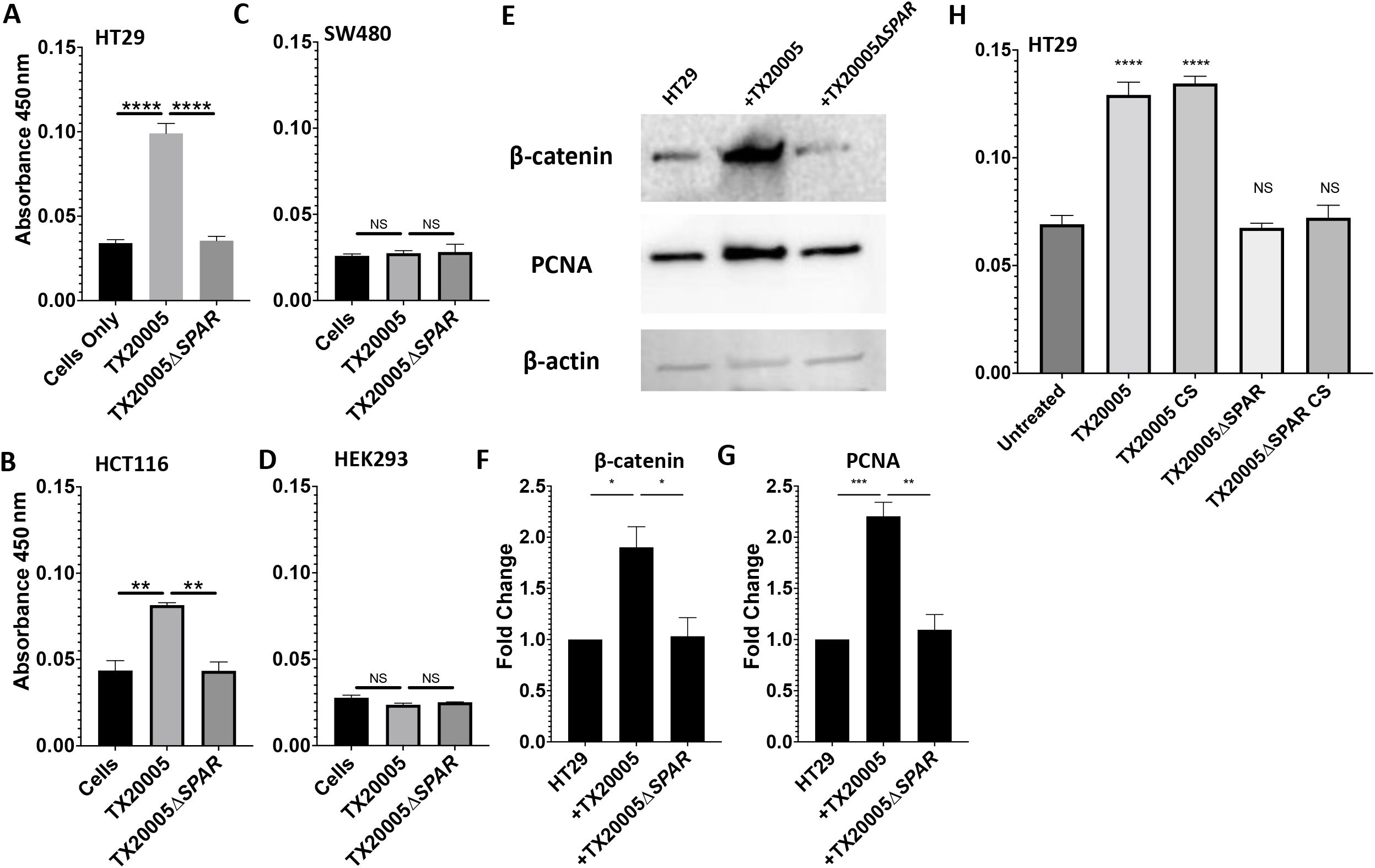
*SPAR* is essential for *Sgg* to stimulate CRC cell proliferation. **A-D**. **Cell proliferation assay.** HT29 (**A**), HCT116 (**B**), SW480 (**C**), and HEK293 (**D**) cells were incubated in the presence or absence of bacteria (MOI=1) or CSs for 24 hours, as described in the Materials and Methods section. Cell proliferation was measured using a CCK-8 kit. Cell-free wells containing only the culture media were served as a blank control to which the absorbance values were normalized. **E-G. Western blot**. Cell lysates from HT29 cells co-cultured in the presence or absence of bacteria (MOI=1) were analyzed by western blot, probed with antibodies against β-catenin, PCNA, and β-actin (**E**). Band intensity was quantified and normalized to β-actin. Fold change in β-catenin (**F**) and PCNA (**G**) were calculated against cells incubated in media only. Data was combined from three biological replicates. **H. Cell proliferation in the presence of *Sgg* CSs.** HT29 cells were cultured in the presence or absence of WT or mutant *Sgg* (MOI=1) or CSs from WT or mutant *Sgg* for 24 hours. Cell proliferation was measured by using a CCK-8 kit. Results were combined from at least three biological replicates. Unpaired two-tailed *t* tests were used for the comparisons. *, *p*<0.05; **, *p*<0.01; ***, *p*<0.001; ****, *p*<0.0001; ns, not significant. Significance in panel H indicates comparison to untreated cells.

To further validate the results, we investigated the effect of the mutant on cell proliferation markers β-catenin and proliferating cell nuclear antigen (PCNA) in HT29 cells (Fig. 3E-G). Cells co-cultured with TX20005 exhibited a significant increase in the level of both β-catenin and PCNA compared to cells cultured in media only, as expected. In contrast, cells co-cultured with the *SPAR* mutant showed no significant increase in the level of either marker compared to cells cultured in media only, further confirming that the *SPAR* locus is essential for *Sgg* to stimulate cell proliferation.

Previous work demonstrated that filter sterilized CSs from WT *Sgg* strains induce CRC cell proliferation (3, 28). Therefore, we examined the effect of CS from TX20005Δ*SPAR* on cell proliferation (Fig. 2H). HT29 cells treated with the WT CS exhibited a significant increase in cell proliferation, as expected, whereas cells treated with the mutant CS did not exhibit any significant increase in cell proliferation. These results suggest that the defect in the *SPAR* mutant to stimulate host cell proliferation is due, at least in part, to the absence of certain secreted factors in the CS. Taken together, these results indicate that *SPAR*, and particularly secreted factors dependent on *SPAR*, is required for *Sgg* to stimulate CRC cell proliferation *in vitro*.

### *Sgg* gut colonization is impaired by *SPAR* deletion

The ability of *Sgg* to colonize the host gut is an important aspect of its pathogenic potential (2). We sought to determine if the *SPAR* locus is involved in *Sgg* gut colonization. Mice were orally gavaged with WT or the *SPAR* deletion mutant. Colon tissues as well as fecal pellets were collected at day 1, 3, and 7 post-bacterial gavage to determine the bacterial load in the samples. At day 1, the *Sgg* bacterial load in the colonic tissues from mice gavage with the WT and the mutant did not significantly differ. At day 3 and 7, the bacterial load of the mutant was significantly decreased compared to that of the WT (Fig. 4A). In the fecal pellets, a similar patten was observed, such that at day 1, the bacterial load of mutant and WT strains did not significantly differ, while At days 3 and 7, the bacterial load of the mutant was decreased significantly compared to that of the WT (Fig. 4B). Taken together, these results indicate that the *SPAR* locus contributes to the colonization capacity of *Sgg* in the normal colon.

**Fig 4.**
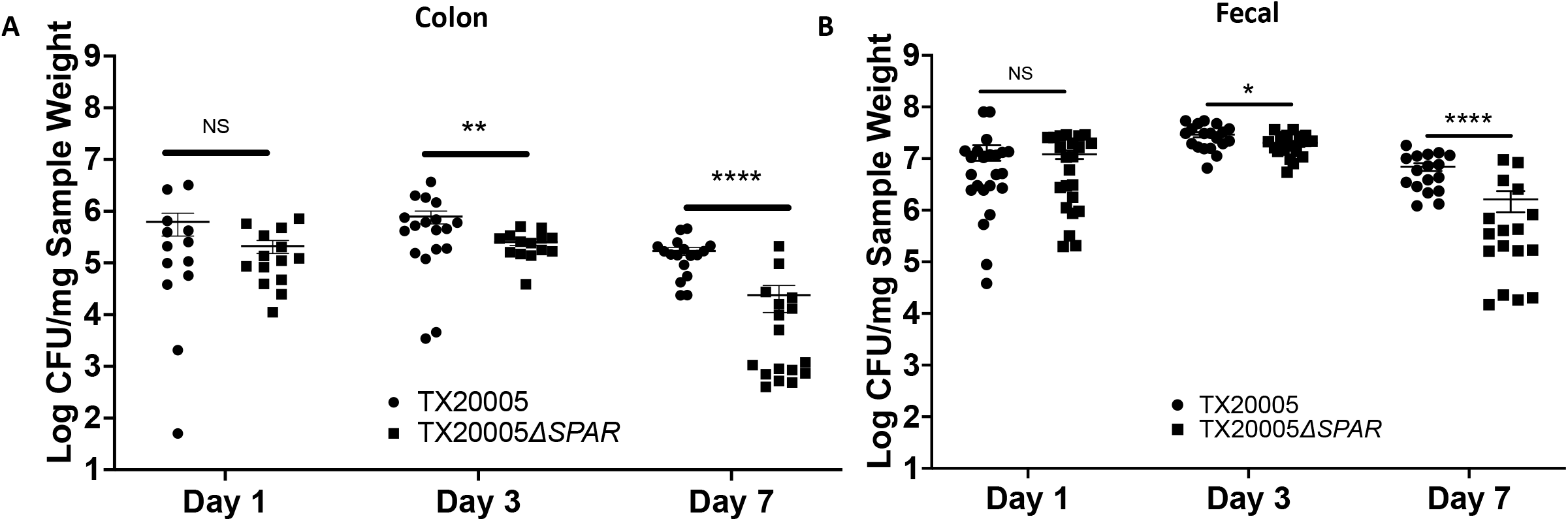
*SPAR* deletion reduced the colonization capacity of *Sgg*. This was performed as described in the Materials and Methods section. Colon tissues (**A**) and fecal pellets (**B**) were weighed, homogenized in sterile PBS and dilution plated onto ESA plates. Mann-Whitney test was used for the comparisons. *, *p*<0.05; **, *p*<0.01; ****, *p*<0.0001; NS, not significant.

### *SPAR* is essential for the ability of *Sgg* to promote the development of colon tumors

Next, we investigated if *SPAR* is important for *Sgg* to promote the development of colon tumors using an AOM-induced CRC model (Fig. 5A). Mice gavaged with the WT strain exhibited a significant increase in the colon tumor burden (Fig. 5B) compared to saline-gavaged mice, consistent with previous results (3, 4). In contrast, mice exposed to the *SPAR* mutant showed no increase in the tumor burden compared to the saline control and a significant reduction compared to the WT-treated mice, suggesting that *SPAR* is important for *Sgg* to promote colon tumors. We also determined the bacterial load in the fecal pellets isolated at the end of the experiment and found no difference in the *Sgg* burden between WT or mutant-treated groups (Fig. 5C). We note that this result could be due to the repeated gavages with bacteria during the experiment and does not necessarily reflect the ability of the strains to colonize tumor-bearing colons in this model. These results together suggest that *SPAR* plays a critical functional role in promoting the development of colon tumors.

**Fig 5.**
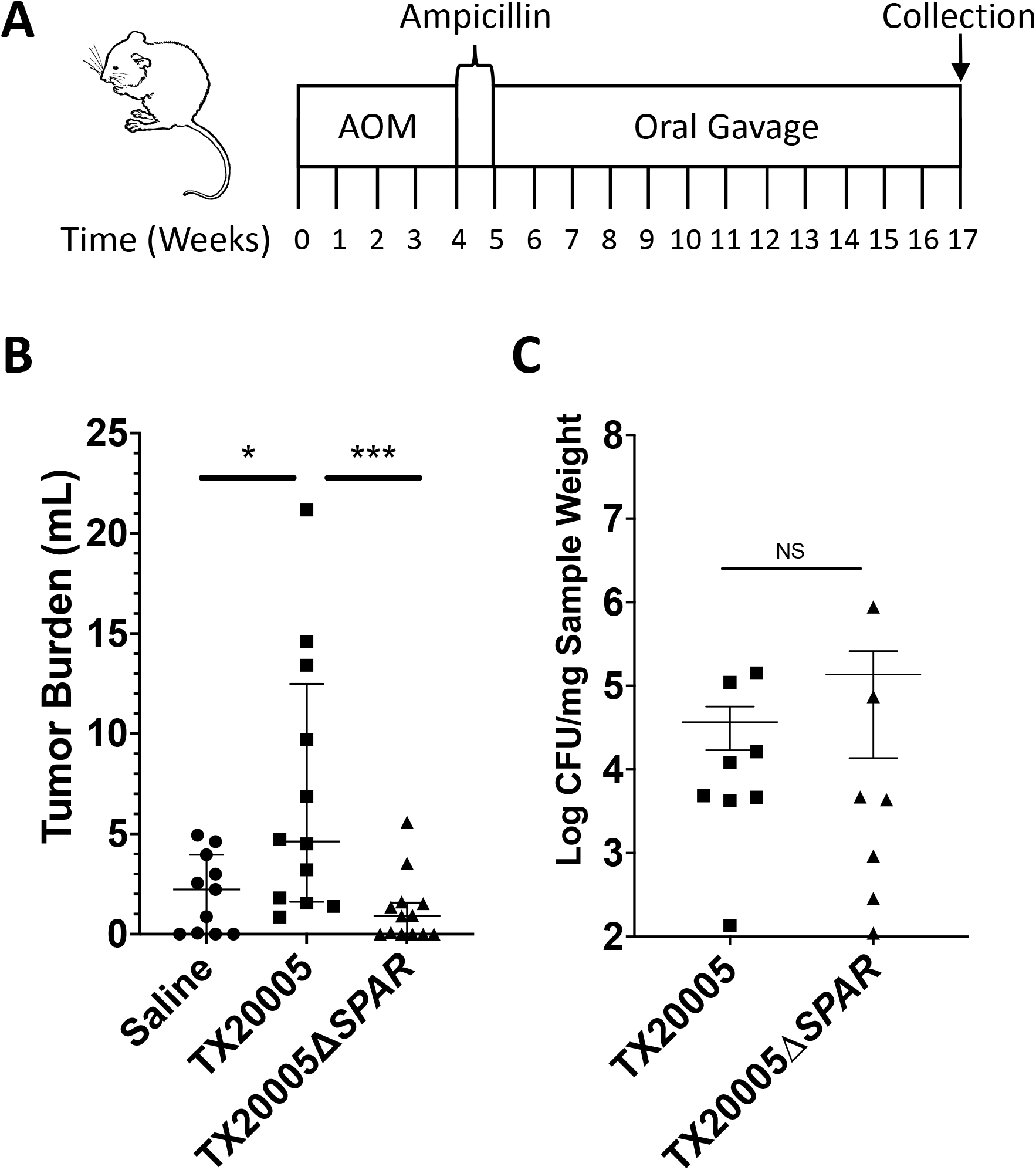
*SPAR* is critical for *Sgg* to promote the development of colon tumors. This was performed as described in the Materials and Methods section. The procedure is illustrated in **A. B.** The number and size of macroscopic tumors were recorded by blinded observers. Tumor burden is calculated as the sum of tumor volumes per mouse. **C.** *Sgg* load in the fecal pellets collected at the experimental endpoint was determined by dilution plating of homogenized pellets onto ESA plates. Mann-Whitney test was used for all comparisons. *, *p*<0.05; **, *p*<0.01; ***, *p*<0.001; ****, *p*, <0.0001; NS, not significant.

## Discussion

In this study, we describe an *Sgg* chromosomal locus, *SPAR*, that plays a key role in *Sgg* pathogenesis. Deletion of *SPAR* led to striking phenotypes *in vitro* and *in vivo*, including a reduced capacity to adhere to CRC cells, decreased colonization of the normal colon, and a complete loss of the ability to stimulate CRC cell proliferation *in vitro* and to promote the development of colon tumors *in vivo*. These results highlight *SPAR* as a critical pathogenicity determinant of *Sgg*.

Our data showed that deletion of the *SPAR* locus resulted in significantly reduced adherence of *Sgg* to HT29 cells. Previous studies showed that T7SS-secreted factors enhance *Sgg* adherence (3). Interestingly, our results revealed that deletion of *SPAR* eliminated the adherence-enhancing activity in the CS, suggesting that the deletion might have affected the T7-secreted factors involved in adherence. In terms of cell proliferation, we showed that the *SPAR* locus is essential for *Sgg* to stimulate certain CRC cell proliferation. Furthermore, the results indicate that *SPAR* is also important for the production of secreted factors responsible for stimulating host cell proliferation. In this aspect, the *SPAR* mutant exhibits a very similar phenotype as that described for a T7SS defective mutant of *Sgg* (TX20005*Δesx*) with respect to adherence and stimulation of cell proliferation (3). One possible reason for the phenotypic similarity is that the *SPAR* locus encodes several putative T7SS effectors (SparG to SparJ). Examination of the TX20005 genome reveals that the previously reported T7SS locus (*Sgg*T7SS^T05^) (3) is the only locus that encodes a set of proteins comprising the T7SS secretion apparatus. Thus, SparG to SparJ are likely secreted by *Sgg*T7SS^T05^. This would imply that one or more of the *SPAR*-encoded putative T7SS effectors are involved in the adherence and the pro-proliferative activity of *Sgg*. On the other hand, the *sparF* gene is predicted to encode a GntR family transcriptional regulator (29). A recent publication indicated that a GntR family transcriptional regulator (OG1RF_11099) of *Enterococcus faecalis* controls the expression of T7SS genes (30). Using Global Align (NCBI), we found that SparF is highly homologous to OG1RF_11099, showing an overall 74% similarity at the amino acid sequence level. Hence, a second possible reason for the phenotypic similarity between *TX20005Δspar* and *TX20005Δesx* is that SparF regulates the expression of genes in the *Sgg*T7SS^T05^ locus.

In a gut colonization model, we observed that deletion of *SPAR* resulted in reduced *Sgg* bacterial load in the normal colon. This could be due to the involvement of *SPAR* in *Sgg* adherence to the colonic epithelium. Additionally, SparJ is a putative T7SS-secreted polymorphic toxin. This family of toxins are often involved in interbacterial competition (26). Thus, SPAR may contribute to *in vivo* colonization by antagonizing other gut commensal bacteria through SparJ.

In the AOM model of CRC, we observed a striking difference between the *SPAR* mutant and the WT bacteria in promoting the development of colon tumors, such that the mutant has completely lost the ability to promote the development of colon tumors. This phenotype is much stronger than that previously reported for the T7SS mutant (3), raising the possibility that in addition to T7SS genes, SparF also regulates the expression of other genes that contribute tumor development *in vivo*. Studies to further investigate the biological activities of SparF, and SparG to SparL are needed to delineate the specific contribution of the *SPAR* locus to the observed phenotypes.

It is also possible that other genes within the *SPAR* locus, SparA-SparE, also play a role, however, their predicted function based on homology search does not provide clues regarding their respective contribution. SparA-C are putative hypothetical proteins. SparD is homologous to OLD family endonucleases. OLD family endonucleases are widely present among bacteria and archaea (31). While their biological function is not completely understood, they contain a C-terminal ATPase domain, as well as an N-terminal Toprim domain, which is believed to be important in DNA replication, recombination, and repair (32, 33). SparE is homologous to GIY-YIG endonuclease (34–38) and PcrA/UvrD helicase. The PcrA/UvrD helicase is an essential helicase in gram-positive bacteria and has been shown to play a role in DNA repair, as well as the replication of small drug-resistance plasmids in *Staphylococcus aureus* and *Bacillus subtilis* (39–45).

In summary, we have identified a chromosomal locus in *Sgg*, *SPAR*, that is critical to the pathogenicity of *Sgg*. We report that deletion of *SPAR* significantly reduces the capacity of *Sgg* to adhere to CRC cells and to colonize the gut. Furthermore, deletion of *SPAR* abrogates the ability for *Sgg* to stimulate CRC cell proliferation and to accelerate colon tumor development. Examination of genes with the locus highlighted several potential candidates responsible for the observed phenotypes. The results suggest a connection between the *SPAR* locus and the T7SS of *Sgg* in terms of additional T7SS effectors encoded by *SPAR* genes and a potential regulator of T7SS expression. Further investigations into the activities of specific *SPAR* proteins will be important for dissecting the intricate *Sgg* pathogenic mechanisms and will open up the path ahead to identify biomarker candidates and targets for clinical prevention and intervention strategies.

## Acknowledgements

This study was supported by funding from the Hamill Foundation, the Cancer Prevention Research Institute of Texas (RP170653) and NIAID (1R21AI151914) to Y. Xu and (T32AI055449-16) to J. Taylor. The content of this manuscript is also available at bioRxiv.org (https://doi.org/10.1101/2022.03.15.484266).

---

**Supplemental Figure S1:**
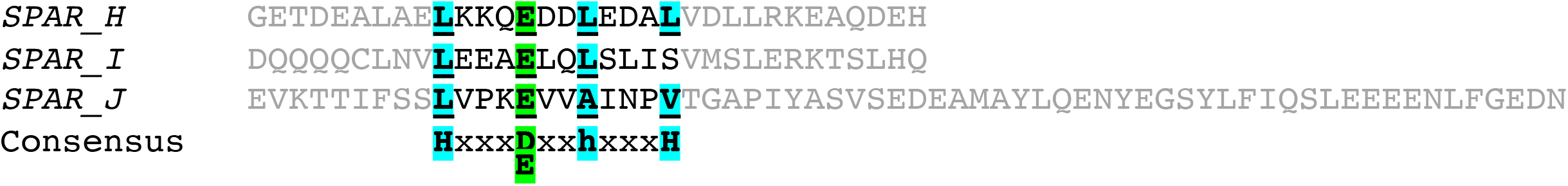
The conserved T7SS sequence in SparH-J. H denotes highly conserved and h less conserved hydrophobic residues.

